# Neurodegeneration-associated mutant TREM2 proteins abortively cycle between the ER and ER–Golgi intermediate compartment

**DOI:** 10.1101/154997

**Authors:** Daniel W. Sirkis, Renan E. Aparicio, Randy Schekman

**Affiliations:** Department of Molecular and Cell Biology, Howard Hughes Medical Institute, University of California, Berkeley, Berkeley, CA 94720

## Abstract

Triggering receptor expressed on myeloid cells 2 (TREM2) is a transmembrane protein expressed on microglia within the brain. Several rare mutations in *TREM2* cause an early-onset form of neurodegeneration when inherited homozygously. Here we investigate how these mutations affect the intracellular transport of TREM2. We find that most pathogenic TREM2 mutant proteins fail to undergo normal maturation in the Golgi complex and show markedly reduced cell surface expression. Prior research has suggested that two such mutants are retained in the endoplasmic reticulum (ER), but we find, using a cell-free COPII vesicle budding reaction, that mutant TREM2 is exported efficiently from the ER. In addition, mutant TREM2 becomes sensitive to cleavage by endoglycosidase D under conditions that inhibit recycling to the ER, indicating that it normally reaches a post-ER compartment. Maturation-defective TREM2 mutants are also efficiently bound by a lectin that recognizes *O*-glycans added in the ER–Golgi intermediate compartment (ERGIC) and *cis* Golgi cisterna. Finally, mutant TREM2 accumulates in the ERGIC in cells depleted of COPI. These results indicate that efficient ER export is not sufficient to enable normal cell surface expression of TREM2. Moreover, our findings suggest that the ERGIC may play an underappreciated role as a quality-control center for mutant and/or malformed membrane proteins.

## INTRODUCTION

Quality control within the eukaryotic secretory pathway occurs primarily at the level of the endoplasmic reticulum (ER). As a general rule, secreted and membrane proteins are screened for proper folding and assembly before gaining competence for export from the ER via COPII vesicles (Geva and Schuldiner, 2014; Barlowe and Helenius, 2016). Proteins that are unable to properly fold within the ER are ultimately degraded via a pathway termed ER-associated degradation, which involves dislocation of the protein from the ER to the cytoplasm and degradation via the proteasome (Vembar and Brodsky, 2008; Brodsky, 2012). However, a small number of malformed transmembrane proteins have been shown to undergo ER export and delivery to the Golgi, followed by retrieval back to the ER (Geva and Schuldiner, 2014; Barlowe and Helenius, 2016). The cases documented thus far generally involve transmembrane subunits of heteromeric protein complexes that are unable to assemble properly. For example, when a single transmembrane subunit (Fet3) of the heterodimeric yeast iron transporter is expressed in the absence of the other subunit, it is transported to an early Golgi cisterna, where it is efficiently returned to the ER by Rer1-mediated retrieval (Sato *et al.*, 2004).

In addition to the few documented cases of transmembrane protein recycling as a means of quality control, a small number of misfolded soluble secretory proteins have been shown to reach the *trans*-Golgi, where they are targeted to the vacuole (lysosome) for degradation (Hong *et al.*, 1996; Coughlan *et al.*, 2004; Kruse *et al.*, 2006; Gelling *et al.*, 2012). In several of these cases, targeting to the vacuole appears to be mediated by the transmembrane receptor, Vps10. Collectively, these observations have suggested that both early and late Golgi cisternae possess quality-control mechanisms that promote the degradation of misfolded or misassembled proteins, either by routing such proteins back to the ER or out to the lysosome for degradation. However, in contrast to the established role of the ER, the extent to which the ERGIC and Golgi participate in protein quality control remains unknown.

Triggering receptor expressed on myeloid cells 2 (TREM2) is a cell surface receptor of the innate immune system that is expressed on dendritic cells, macrophages, osteclasts and microglia (Bouchon *et al.*, 2001; Colonna, 2003; Turnbull *et al.*, 2006; Butovsky *et al.*, 2014). Homozygous and compound heterozygous mutations in the *TREM2* gene have been linked to a syndromic form of early-onset frontotemporal dementia (FTD) that usually occurs with recurrent bone fractures (Nasu Hakola disease [NHD]; also known as polycystic lipomembranous osteodysplasia with sclerosing leukoencephalopathy [PLOSL]) (Paloneva *et al.*, 2002; Klünemann *et al.*, 2005). More recently, a rare variant in the *TREM2* gene (R47H) has been associated with a large increase in the risk of developing late-onset Alzheimer’s disease (Jonsson *et al.*, 2013; Guerreiro *et al.*, 2013a). Recent work has focused on how the R47H variant alters the biochemical properties of TREM2 (Wang *et al.*, 2015; Kober *et al.*, 2016; Yeh *et al.*, 2016), and evidence is emerging that this variant is associated with impaired ligand binding.

On the other hand, TREM2 missense mutants Y38C and T66M, which cause early-onset neurodegeneration, have been shown to result in reduced protein maturation and impaired cell surface expression (Kleinberger *et al.*, 2014; Park *et al.*, 2015). In addition, the steady-state localization of these mutants appears to be restricted to the ER (Kleinberger *et al.*, 2014; Park *et al.*, 2015). Recent biochemical analyses have also indicated that these mutations impair TREM2’s ability to bind ligand (Yeh *et al.*, 2016) and may also promote misfolding of the lumenal Ig-like domain to which these mutations map (Kober *et al.*, 2016).

In the course of studying the intracellular transport of TREM2 mutants within the secretory pathway, we made the surprising discovery that multiple pathogenic TREM2 missense mutants previously presumed to be retained in the ER are in fact competent for efficient ER export and delivery to the ERGIC, from which they appear to be actively retrieved to the ER.

## RESULTS

TREM2 disease mutants Y38C and T66M have been shown by others to have strongly impaired cell surface expression (Kleinberger *et al.*, 2014; Park *et al.*, 2015). Moreover, these mutants were shown to accumulate in an immature form, and immunofluorescence results suggested they might be retained within the ER. To further investigate and expand upon these findings, we began by generating the six known missense mutations (Y38C, T66M, D86V, V126G, D134G and K186N) implicated in NHD or early-onset FTD lacking the bone phenotype (Paloneva *et al.*, 2002; Klünemann *et al.*, 2005; Guerreiro *et al.*, 2013b; 2013c; Le Ber *et al.*, 2014). We transiently expressed HA-tagged versions of the WT and mutant proteins in HEK-293T cells and found by immunoblotting that the four mutations located within the Ig-like domain (Y38C, T66M, D86V, and V126G) resulted in TREM2 accumulating in an apparently immature form (Figure 1A). Mutants Y38C and T66M have previously been shown to be sensitive to cleavage by the endoglycosidase, endo H (Park *et al.*, 2015), suggesting that these mutants are not processed by α-mannosidase II in the *medial* Golgi. We confirmed that, in our expression system, mutant Y38C was sensitive to cleavage by both endo H and endo F1 (Freeze and Kranz, 2006), indicating that it had no complex, *N*-linked glycans (Figure 1B).

**Figure 1.**
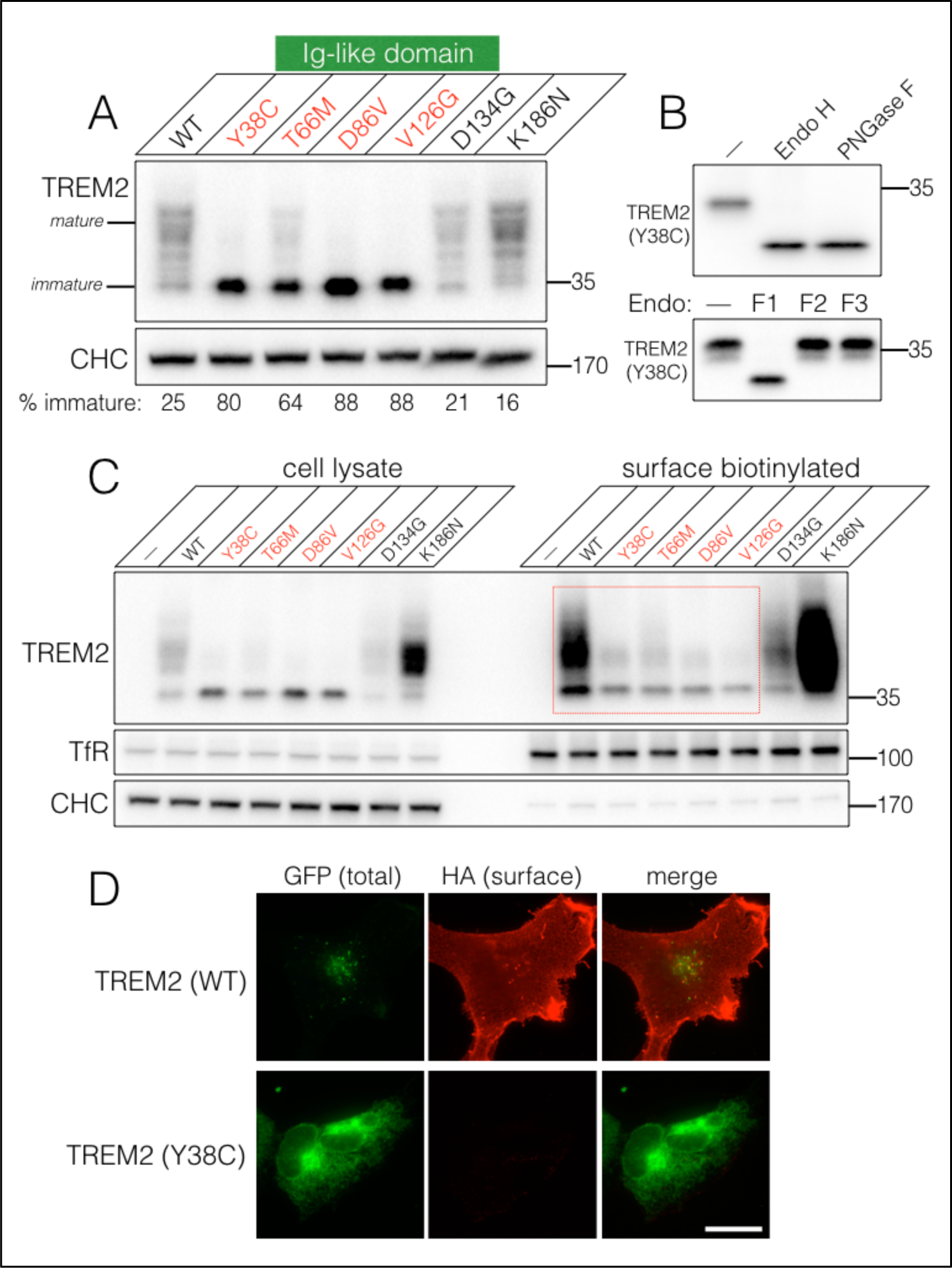
Neurodegeneration-associated TREM2 point mutants display defective maturation and cell surface expression. (A) HEK-293T cells were transfected with WT HA-TREM2 or neurodegeneration-associated point mutants. One day after transfections, the cells were lysed and the lysates analyzed by immunoblotting. Mutants Y38C, T66M, D86V and V126G show defects in protein maturation. Maturation was quantified by taking the ratio of the amount of immature TREM2 (the fastest-migrating band) to total TREM2 signal. CHC was used as loading control. (B) Digests of cell lysates reveal that TREM2 (Y38C) is sensitive to cleavage by endo H, endo F1, and PNGase F, indicating that it contains non-complex *N*-linked glycans. (C) Cell surface biotinylation reveals that the four TREM2 point mutants displaying maturation defects also show marked reductions in cell surface expression. TfR and CHC were used as positive and negative controls, respectively, for biotinylation and pulldown. TfR also served as a loading control. (D) U2OS cells were transiently transfected with WT or Y38C HA-TREM2-GFP and prepared for immunofluorescence microscopy the following day. Labeling of fixed, non-permeabilized cells for HA confirms that TREM2 (Y38C) shows impaired cell surface expression. Scale bar indicates 20 μm.

We next determined the cell-surface expression phenotypes of the six mutants using surface biotinylation, and found that the same four Ig-like domain mutants that accumulated in an immature form also showed large defects in cell-surface expression (Figure 1C). Mutant D134G showed lower overall expression but no maturation or surface expression defect, while mutant K186N accumulated at high levels on the cell surface (Figure 1C). The latter mutation is presumed to render TREM2 defective for signal transduction due to the loss of a conserved transmembrane lysine residue that is required for association with the signaling adaptor protein DAP12 (Bouchon *et al.*, 2000; Call *et al.*, 2010). We further assessed the cell-surface expression of mutant Y38C using immunofluorescence analysis. In particular, we labeled the cell-surface pool of TREM2 in fixed cells using a monoclonal antibody (mAb) directed against an extracellular HA tag inserted downstream of TREM2’s signal peptide, and found robust surface expression for WT TREM2 but very weak surface expression for mutant Y38C (Figure 1D). A cytoplasmic GFP fusion on TREM2 showed that both proteins were robustly expressed within the cell. However, as reported by others (Kleinberger *et al.*, 2014; Park *et al.*, 2015), we found that mutant Y38C, in contrast to the WT protein, showed a striking reticular, ER-like staining pattern.

To directly test the hypothesis that the maturation-defective TREM2 disease mutants were retained within the ER, we performed cell-free COPII vesicle budding reactions. We transiently expressed HA-tagged WT or Y38C TREM2 in HEK-293T cells and prepared permeabilized cells for the budding assay (Wilson *et al.*, 1995; Merte *et al.*, 2010). Membranes expressing WT TREM2 showed cytosol-dependent budding of TREM2 that was inhibited by the addition of either recombinant Sar1A (H79G) or the non-hydrolyzable GTP analog GTPγS, which prevent COPII vesicle formation at the ER (Figure 2A). Similarly but quite unexpectedly, we found that membranes expressing TREM2 (Y38C) showed robust budding of the mutant protein, suggesting that mutant Y38C was not trapped in the ER (Figure 2A). We next performed several control experiments to determine whether our budding assay was capable of detecting ER retention of a synthetic, misfolded version of TREM2. To generate such a form of TREM2, we first treated cells expressing TREM2 (Y38C) with tunicamycin for 4 hr prior to generating permeabilized cells for the budding assay. This treatment duration enabled the accumulation of an unglycosylated form of mutant TREM2 (running at ~25 kD) in roughly equal proportion to the glycosylated form (Figure 2B, see input lane). Vesicle budding reactions performed with these membranes showed that unglycosylated TREM2 (Y38C) was exported poorly from the ER, whereas glycosylated TREM2 (Y38C), derived from either control or treated membranes, showed normal budding efficiency (Figure 2B). We independently confirmed these findings by mutating the two predicted *N*-linked glycosylation sites within TREM2. As expected, the non-glycosylatable form of TREM2 (Y38C) showed greatly diminished budding capacity compared to glycosylated Y38C (Figure 2C). Thus, TREM2 (Y38C) does not appear to be grossly misfolded and is exported from the ER as efficiently as WT TREM2.

**Figure 2.**
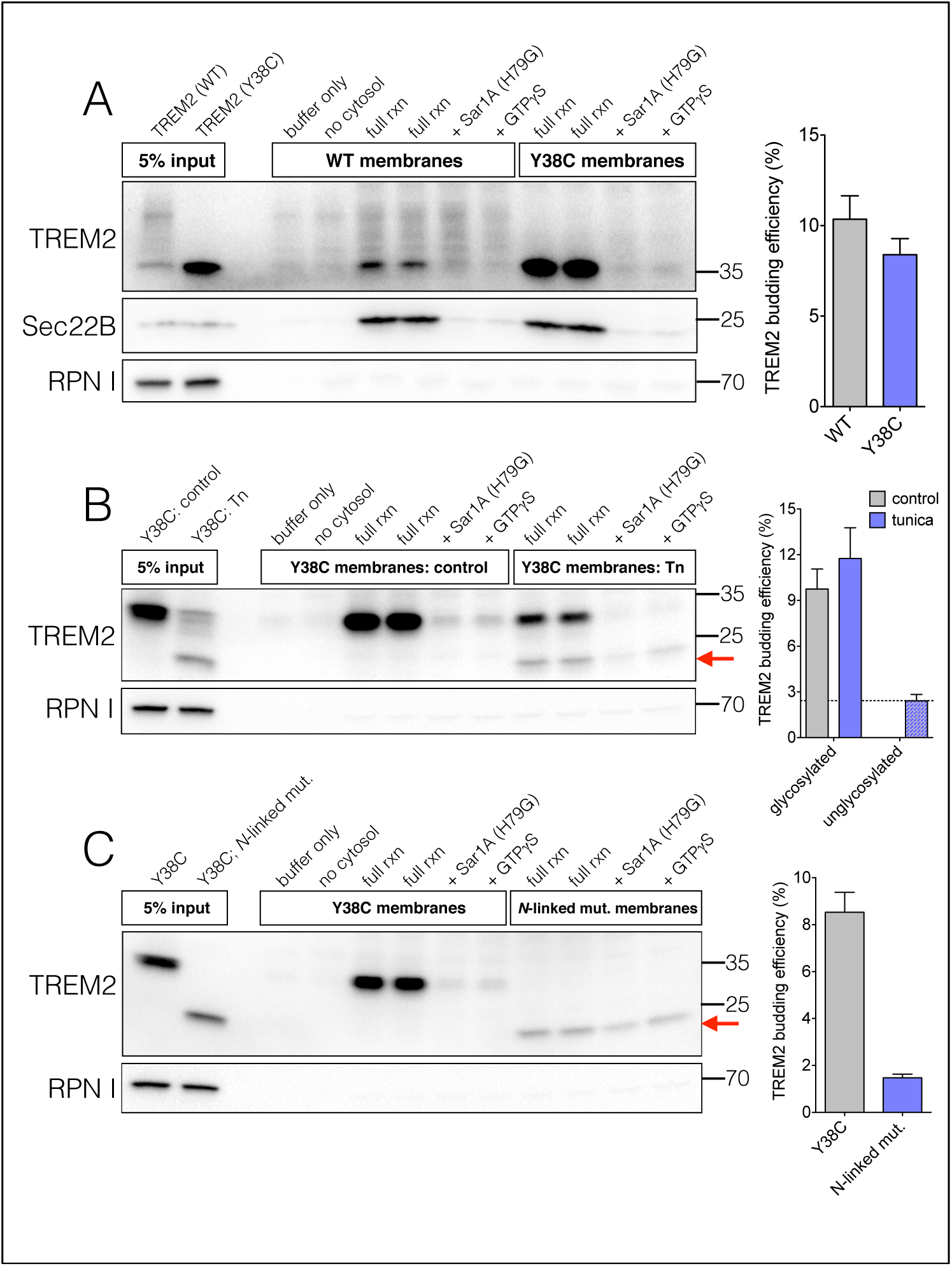
TREM2 (Y38C) is competent for efficient ER export. (A) HEK-293T cells were transfected with HA-tagged WT or Y38C TREM2. One day after transfection, cell membranes were prepared for the COPII vesicle budding assay. The budding reaction was carried out at 30°C for 30 min and vesicles were isolated by differential centrifugation. Immunoblotting of the vesicle fraction reveals that both WT and Y38C TREM2 show cytosol-dependent budding that is sensitive to the addition of recombinant Sar1A (H79G) and GTPγS. Budding efficiency (right), calculated by comparing signals from budded lanes to the respective 5% membrane input lanes, indicates that WT and Y38C TREM2 are exported from the ER with similar efficiency. Sec22B serves as a positive control for ER budding and RPNI, an ER-resident protein, serves as a negative control. (B) Pretreatment of cells with tunicamycin (Tn) for 4 h prior to harvesting leads to accumulation of a non-glycosylated form of TREM2 (Y38C). This form buds poorly from the ER relative to the glycosylated form, demonstrating the specificity of budding in this assay. (C) A version of TREM2 (Y38C) in which both predicted *N*-linked glycosylation sites are abolished buds inefficiently from the ER. Red arrows indicate the location of non-glycosylated form of TREM2 (Y38C) in (B and C). The plotted budding efficiency values (A-C, right) are derived from 2-3 independent experiments and error bars indicate SEM.

The efficient ER budding observed for TREM2 (Y38C) was unlikely to be an artifact of the cell type used or of transient expression, as we observed similarly non-defective budding behavior for mutant Y38C in membranes derived from murine N9 microglial cells stably expressing low levels of HA-TREM2 (Supplemental Figure S1A) and HATREM2-GFP (Supplemental Figure S1B). In addition, the budding behavior observed in HEK-293T membranes was not unique to mutant Y38C, as we also observed efficient budding for mutants T66M and V126G, which displayed comparable defects in protein maturation and cell-surface expression (Supplemental Figure S1, C and D).

The results obtained thus far suggest that, despite having a maturation defect and showing very poor surface expression, the TREM2 disease mutants are not retained within the ER. To determine whether these mutants were capable of reaching post-ER compartments *in vivo*, we devised two biochemical assays and one morphological assay. We first made use of the endoglycosidase, endo D, which reports on the delivery of glycoproteins to the ERGIC (Alvarez *et al.*, 1999) or *cis*-Golgi (Beckers *et al.*, 1987) (Figure 3A). Normally, glycoproteins transiting the secretory pathway display sensitivity to endo D only transiently – after encountering α-mannosidase I in the ERGIC or *cis* Golgi cisterna but before encountering GlcNAc transferase I in the *medial* Golgi cisterna (Freeze and Kranz, 2006). However, by using the CHO-Lec1 cell line, which is deficient in GlcNAc transferase I activity, endo D sensitivity is retained for glycoproteins at any point after being acted on by α-mannosidase I (Beckers *et al.*, 1987; Freeze and Kranz, 2006). Before assessing endo D sensitivity, we first confirmed that TREM2 (Y38C) showed a robust cell-surface expression defect in CHO-Lec1 cells (Supplemental Figure S2), indicating that this cell line was suitable for studying the trafficking of mutant TREM2.

**Figure 3.**
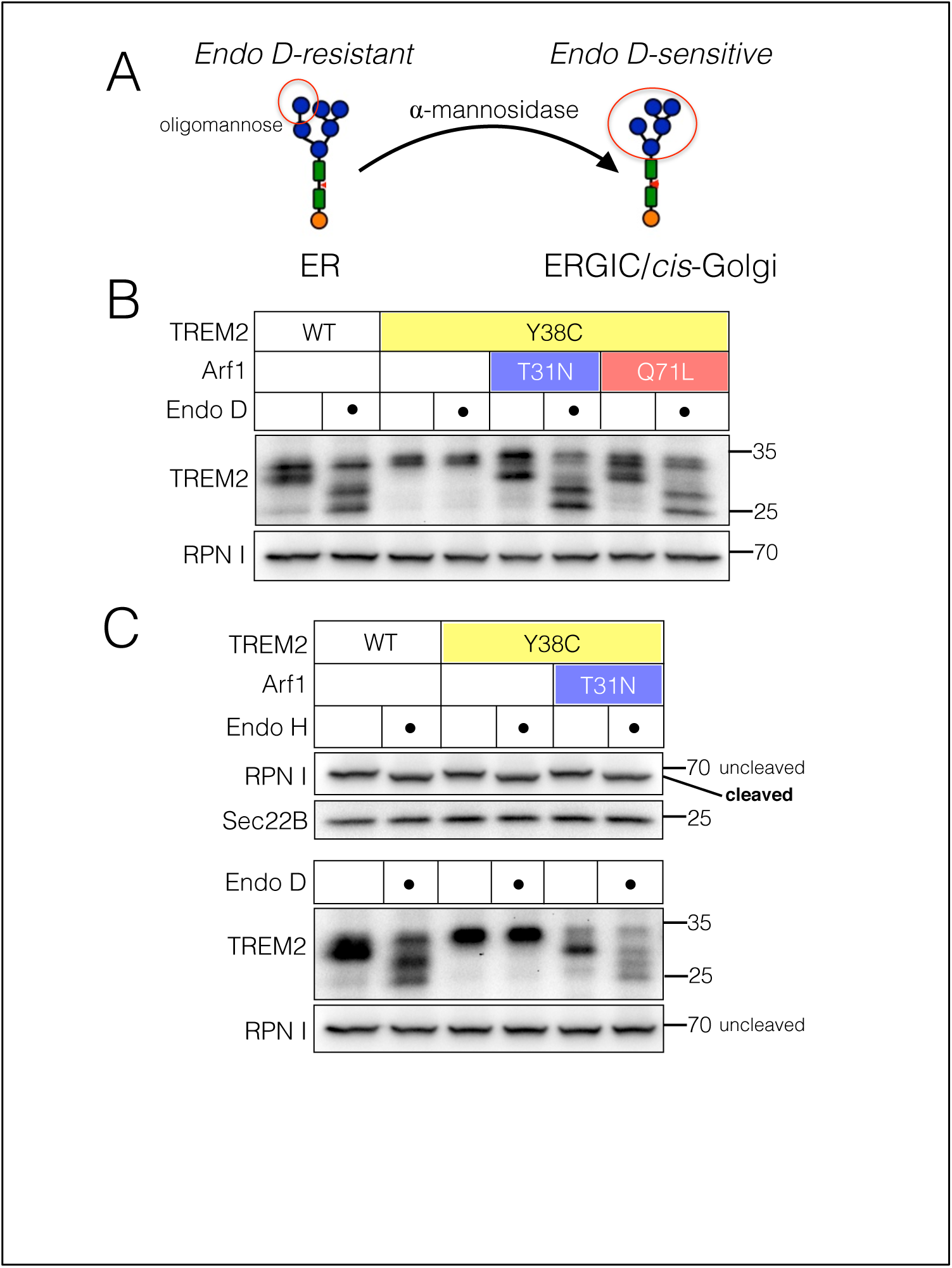
Mutant TREM2 gains access to a post-ER compartment *in vivo*. (A) Schematic illustrating the generation of a five-mannose-containing *N*-linked glycan by α- mannosidase-I in the ERGIC or *cis*-Golgi to produce a protein that is sensitive to cleavage by endo D. (B) CHO-Lec1 cells were co-transfected with either WT or Y38C HA-TREM2 plus either Arf1 (T31N), Arf1 (Q71L) or empty vector. Cells were harvested one day after transfection and the lysates subjected to digest by endo D. WT TREM2 is sensitive to endo D, while mutant Y38C is normally insensitive. Expression of either Arf1 (T31N) or (Q71L) uncovers endo D sensitivity for TREM2 (Y38C). RPN I serves as a loading control. (C) The ER-resident protein RPN I remains insensitive to endo D under conditions that render TREM2 (Y38C) sensitive to endo D. RPN I cleavage by endo H is detectable under the same experimental conditions. Sec22B serves as a loading control.

As expected for a cell surface protein, we found that WT TREM2 was sensitive to cleavage by endo D (Figure 3B). Conversely, we found that mutant Y38C was fully resistant to endo D, raising the possibility that this mutant was unable to reach the ERGIC or *cis* Golgi. However, another possibility, consistent with the budding assay results that demonstrated robust ER export, was that mutant Y38C was delivered to a post-ER compartment but very efficiently returned to the ER via the COPI-dependent, retrograde transport pathway. To test this possibility, we co-expressed, together with TREM2 (Y38C), either of two nucleotide-locked forms of Arf1 to block the COPI retrieval pathway (Lippincott-Schwartz *et al.*, 1998). Strikingly, co-expression of these Arf1 mutants rendered the Y38C mutant sensitive to endo D (Figure 3B), indicating that Arf1/COPI-mediated recycling of mutant TREM2 from the ERGIC or Golgi to the ER might be responsible for their reported steady-state localization to the ER (Kleinberger *et al.*, 2014; Park *et al.*, 2015). Because α-mannosidase I is the key enzyme mediating sensitivity to cleavage by endo D, we tested the possibility that expression of the nucleotide-locked forms of Arf1 indirectly led to an accumulation of α-mannosidase I in the ER, which would complicate the interpretation of the results. To address this possibility, we tested whether the ER-resident glycoprotein ribophorin I (RPN I) became sensitive to endo D upon mutant Arf1 expression. However, we found that RPN I remained endo D-resistant under all conditions tested (Figure 3C, bottom), despite showing the expected sensitivity to endo H (Figure 3C, top). Moreover, since endo D and H cleave *N*-linked glycans at the same position, our experimental conditions should have been suitable for detecting any potential endo D sensitivity for RPN I. Thus, mutant TREM2 appears capable of reaching the ERGIC or early Golgi.

The output of the above assay (endo D sensitivity) depended on the use of Arf1 mutants. Thus, we developed an independent biochemical assay reporting on the delivery of mutant TREM2 to post-ER compartments that did not involve the use of Arf1 mutants. We reasoned that a sensitive detection system for the acquisition of *O*-linked glycans, which are added in the ERGIC and Golgi apparatus (Dunphy and Rothman, 1985; Krijnse-Locker *et al.*, 1994), might provide an alternative means to assess mutant TREM2’s transport itinerary. We started by adding a synthetic 8X repeat of an *O-*glycosylation motif (Asada *et al.*, 1999; Yoneda *et al.*, 2001) to the lumenal portion of TREM2, just downstream of the N-terminal HA tag. To specifically detect *O*-glycosylated TREM2 species, we prepared cell lysates and performed affinity chromatography using an immobilized form of the *O*-glycosylation-specific lectin, jacalin (Hortin and Trimpe, 1990), which recognizes *O*-linked N-acetylgalactosamine (GalNAc) and galactose (Aucouturier *et al.*, 1987).

We first expressed these TREM2 constructs in CHO-Lec1 cells and found highly efficient pulldown of mutant Y38C by jacalin (Figure 4A). We reasoned that such efficient pulldown might be the result of repeated cycling of mutant TREM2 into and out of the ERGIC or Golgi, which would be consistent with both the budding results and Arf1/COPI-dependent recycling of mutant TREM2 to the ER. As positive and negative control proteins for the pulldown, we used transferrin receptor, which is *O*-glycosylated (Do and Cummings, 1992), and Sec22B, which is not glycosylated (Schuldiner *et al.*, 2008).

**Figure 4.**
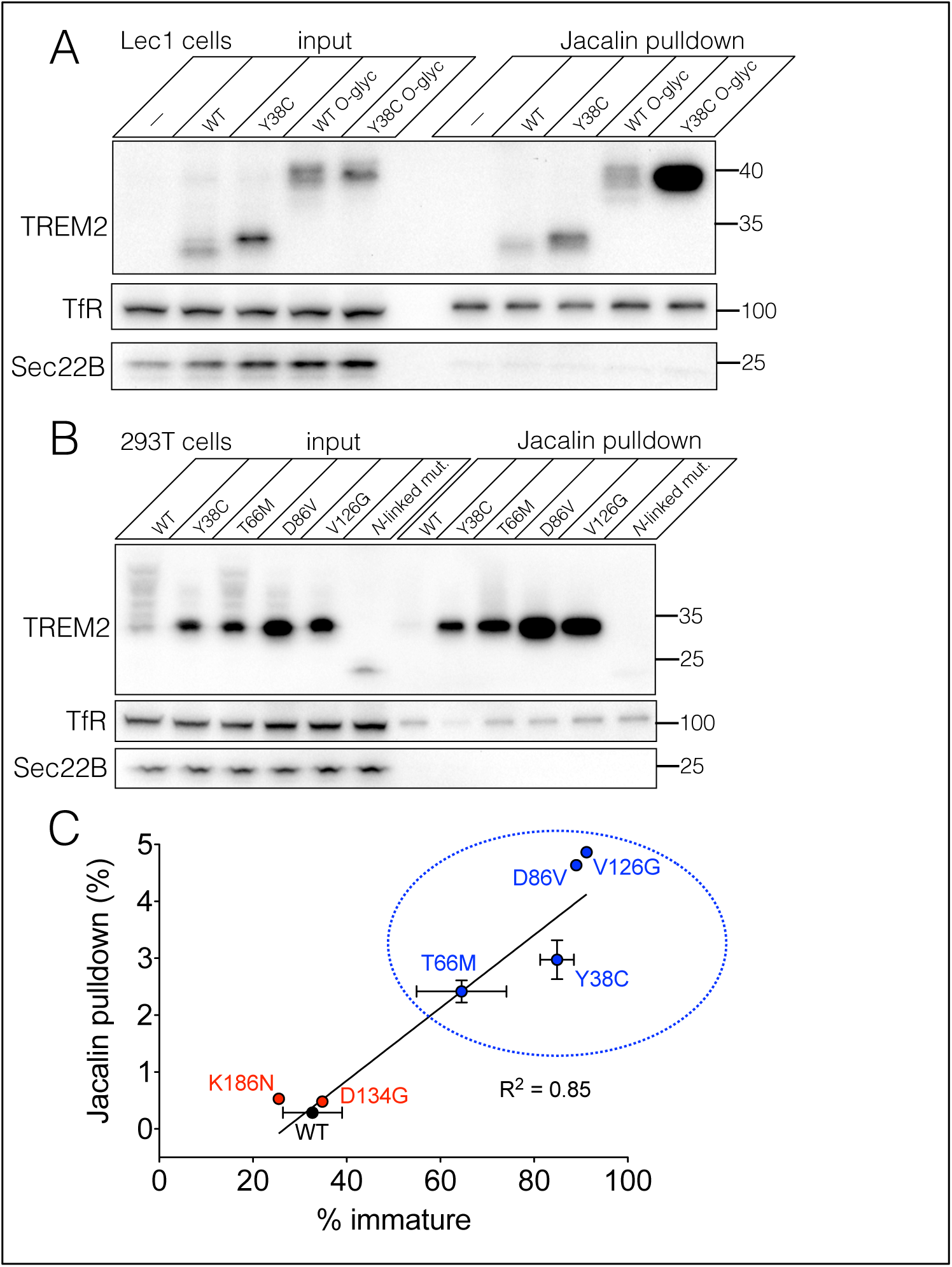
TREM2 point mutants displaying defective maturation and surface expression are modified by *O*-glycosylation. (A) CHO-Lec1 cells were transfected with WT or Y38C HA-TREM2 lacking or containing a synthetic *O*-glycosylation motif. One day after transfection, cells were harvested and the lysates subjected to affinity chromatography with jacalin. TREM2 (Y38C) containing the *O*-glycosylation motif shows robust pulldown by jacalin relative to all other constructs. TfR and Sec22B serve as positive and negative controls, respectively, for pulldown by jacalin. (B) HEK-293T cells were transfected with WT HA-TREM2 or the indicated mutants and processed for jacalin pulldown the following day. Mutants Y38C, T66M, D86V, V126G are all efficiently bound by jacalin while WT TREM2 is not. An ER-retained version of mutant Y38C that harbors mutated *N*-linked glycosylation sites is no longer bound by jacalin. (C) Jacalin pulldown efficiency was quantified by comparing the signal intensity for pulldown lanes to the corresponding input lanes and plotting these values against the corresponding protein maturation values, calculated as described above. Vertical and horizontal error bars (SEM) are displayed for WT, Y38C, and T66M, for which values were obtained from at least three independent experiments. Linear regression indicates a robust correlation between jacalin pulldown efficiency and protein maturation (R^2^ = 0.85). Mutants Y38C, T66M, D86V, and V126G cluster together on the graph (indicated by dashed blue oval).

Because CHO-Lec1 cells have a partially defective *N*-linked glycosylation pathway (Freeze and Kranz, 2006), we sought to confirm the results for Y38C and test additional disease mutants in HEK-293T cells. To our surprise, we found that all four disease mutations within the Ig-like domain resulted in very robust *O*-glycosylation of TREM2, such that it was unnecessary to utilize the synthetic *O*-glycosylation motif that we originally developed for use in CHO-Lec1 cells (Figure 4B). In contrast to the mutant proteins, WT TREM2 showed only very weak pulldown (Figure 4B), indicating that it was not normally *O*-glycosylated.

Because we showed in the *in vitro* budding assay that a version of the Y38C mutant rendered defective for *N*-linked glycosylation was retained in the ER, we used this construct to ask whether the *O*-linked glycosylation we were observing could still occur when a disease mutant was retained in the ER. As expected, we found that the *N*glycosylation-defective version of mutant Y38C was not pulled down by jacalin (Figure 4B), suggesting that the modification recognized by jacalin occurs outside of the ER. In addition, we obtained similar results using tunicamycin treatment of cells expressing mutant TREM2 (Supplemental Figure S3B). We also confirmed that jacalin did not recognize an unusual type of *N*-linked glycosylation, because enzymatic removal of mutant TREM2’s *N*-linked glycans by endo H or PNGase F did not affect the efficiency of pulldown (Supplemental Figure S3C). We also tested the possibility that the observed jacalin binding was due to a cytoplasmic *O*-linked modification (Wells and Hart, 2003). However, we found that a version of the Y38C mutant lacking a cytoplasmic tail was still efficiently bound by jacalin, indicating that the modification occurs lumenally (Supplemental Figure S3A).

Mutants D134G and K186N, which do not display maturation or surface expression defects, were found not to be bound by jacalin (Supplemental Figure S3A). Overall, when the pulldown efficiency of the TREM2 disease mutants was plotted against their relative maturation, we found that the four Ig-like domain mutants clustered together (Figure 4C). Thus, an inability to undergo complex *N*-linked glycosylation shared among the Ig-like domain mutants appears to be associated with an alternative, *O*-linked glycosylation pathway. In addition, the relatively low efficiency of mutant TREM2 pulldown by jacalin (~2-5% of total) is consistent with our inability to detect endo D sensitivity for mutant TREM2 when retrieval to the ER was left intact.

To complement the results of the two independent biochemical assays we used to assess delivery to the ERGIC and/or Golgi *in vivo*, we developed an alternative, immunofluorescence microscopy-based approach that could provide a morphological readout for mutant TREM2’s localization. Because mutant TREM2 has been reported to localize to the ER at steady state (Kleinberger *et al.*, 2014; Park *et al.*, 2015), we reasoned that it would likely be difficult to visualize a sub-population of the protein that transiently passes through the ERGIC or Golgi. Given that the endo D assay suggested that mutant TREM2 is returned to the ER via the COPI pathway, we reasoned that COPI depletion by knockdown might be sufficient to retain mutant TREM2 in a post-ER compartment. For these experiments, we utilized human osteosarcoma U2OS cells which are flat and therefore ideal for immunofluorescence microscopy (Martinez *et al.*, 2013). We first confirmed that in cells transfected with control, non-targeting siRNA, TREM2 (Y38C)-GFP localized to the ER (Figure 5A). In particular, TREM2 (Y38C)-GFP in control cells showed striking colocalization with reticular, calnexin-positive structures and did not colocalize with ERGIC-53 (Figure 5, A and B).

**Figure 5.**
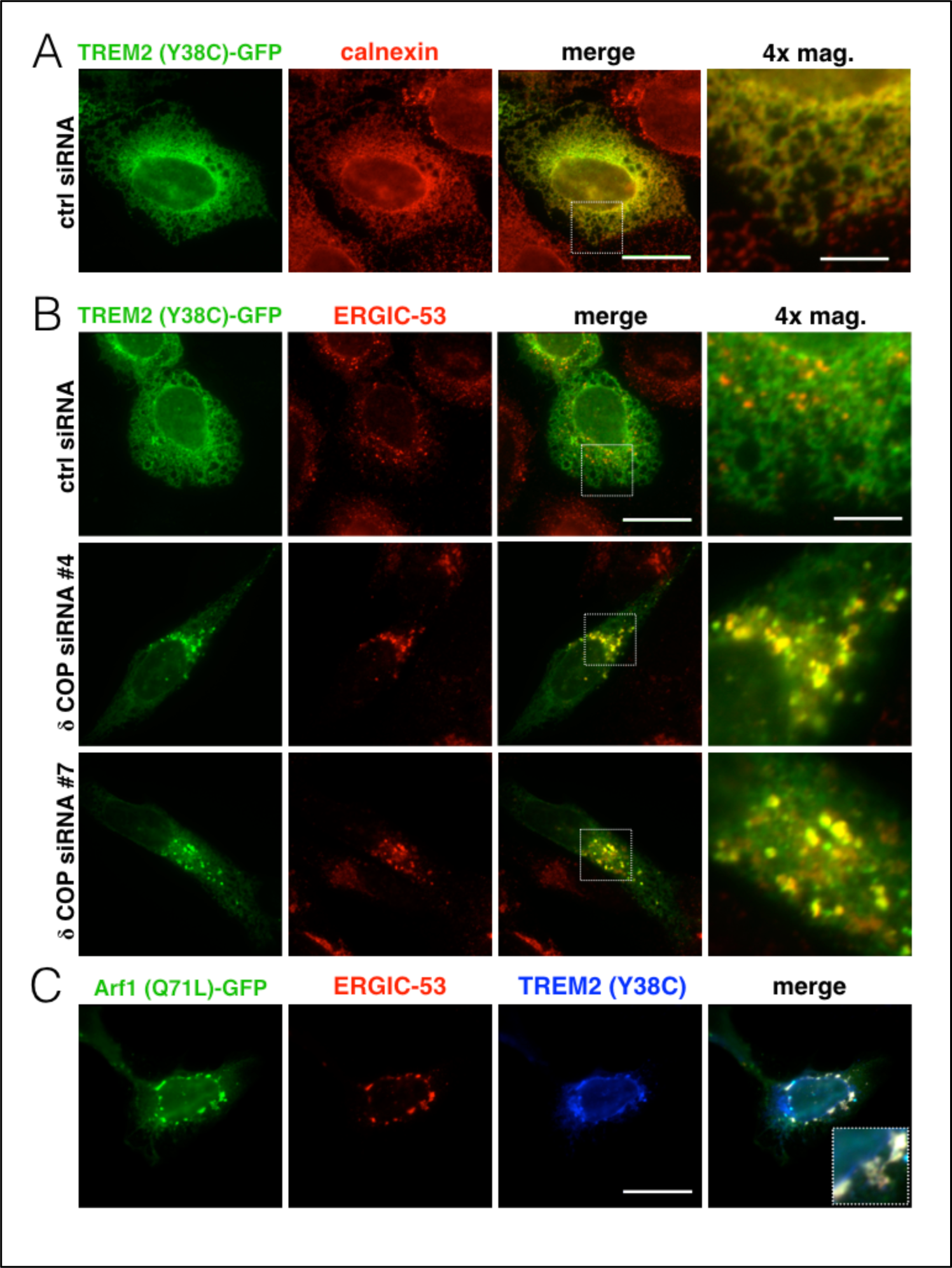
Depletion of COPI results in retention of mutant TREM2 within the ERGIC. (A and B) U2OS cells were transfected with either of two independent siRNAs targeting δ COP or a non-targeting control siRNA, followed by transfection with the indicated plasmid the next day. Two days after the siRNA transfection, cells were processed for microscopy and stained for calnexin or ERGIC-53. (A) TREM2 (Y38C)- GFP shows strong colocalization with calnexin-positive reticular structures in cells transfected with control siRNA. (B, top row) TREM2 (Y38C)-GFP does not colocalize with ERGIC-53 in control cells. (B, middle and bottom rows) TREM2 (Y38C)-GFP shows striking colocalization with large, ERGIC-53-positive puncta after transfection with either of two independent siRNAs targeting δ COP. The results shown are representative of two independent experiments and >30 cells imaged. (C) Co-transfection of cells with Arf1 (Q71L)-GFP and untagged TREM2 (Y38C) results in the appearance of puncta positive for Arf1 (Q71L)-GFP, ERGIC-53 and TREM2 (Y38C). The inset (merged image, bottom right) shows a 3x-magnified detail of triple-positive puncta. Scale bars (A-C) indicate 20 μm for standard magnification and 5 μm for the 4x-magnified images.

To deplete COPI by RNAi, we transfected cells with siRNA targeting coatomer subunit δ (*ARCN1* gene) (Bettayeb *et al.*, 2016). We observed >80% reduction in δ COP levels as assessed by immunoblotting two days post-transfection, and cells showed no obvious signs of toxicity from this level of knockdown. Cells in which δ COP had been depleted by either of two distinct siRNAs showed a striking change in the localization of TREM2 (Y38C), suggesting that the effect observed was unlikely to be off-target. In particular, TREM2 (Y38C)-GFP in COPI-depleted cells accumulated in large, ERGIC-53-positive puncta (Figure 5B, middle and bottom rows). In addition, we could recapitulate the effect of COPI knockdown by expressing the Arf1 (Q71L) mutant. In particular, in cells cotransfected with Arf1 (Q71L)-GFP and untagged TREM2 (Y38C), we observed large, TREM2-positive puncta that were also positive for Arf1 (Q71L) and ERGIC-53 (Figure 5C). Next, to determine whether the TREM2-containing puncta we identified upon COPI knockdown colocalized with markers of other organelles, we also labeled cells for markers of ER, ER exit sites, Golgi, early endosomes and late endosomes/lysosomes (Supplemental Figure S4, A–E). However, we found that the TREM2-positive puncta did not show appreciable colocalization with any of these additional markers. Thus, mutant TREM2 is effectively retained within the ERGIC upon depletion of COPI. Moreover, these results provide morphological corroboration for the biochemical assays used, both of which indicated that mutant TREM2 reaches a post-ER compartment *in vivo*.

## DISCUSSION

Of the few known examples of retrograde recycling of membrane proteins from an early Golgi cisterna to the ER as means of quality control, most involve transmembrane subunits of multimeric complexes expressed in the absence of their obligate binding partner(s). An early discovery of this phenomenon involved recycling of class I major histocompatibility complex (MHC) molecules in a cell line that was defective for MHC assembly (Hsu *et al.*, 1991). In particular, it was found that MHC molecules appearing morphologically to be retained in the ER were in fact capable of actively cycling between the ER and an early Golgi compartment. More recently, it has been demonstrated that class I MHC molecules sub-optimally loaded with antigenic peptide are similarly retrieved to the ER from the *cis* Golgi compartment (Garstka *et al.*, 2007; Howe *et al.*, 2009). In addition, the temperature-sensitive folding mutant of the vesicular stomatitis virus G protein (tsO45 VSV-G) can reach the early Golgi when over-expressed, and appears to recycle to the ER by remaining in complex with the KDEL-motif containing ER chaperone, BiP (Hammond and Helenius, 1994). Along similar lines, when the T-cell receptor (TCR) α chain is expressed heterologously in a cell line lacking the other subunits of the TCR, a fraction of TCRα reaches the Golgi and recycles via BiP- and KDEL receptor-mediated transport (Yamamoto *et al.*, 2001). In addition, when the yeast plasma membrane iron transporter subunit Fet3 is expressed in the absence of the other subunit (Ftr1), it shows a steady-state localization to the ER but depends on Rer1-mediated recycling from the early Golgi to maintain this localization (Sato *et al.*, 2004). A common theme among these examples is the role of transmembrane receptors and the COPI pathway acting in concert to promote quality control in the early secretory pathway. In the case of Rer1-mediated recycling of Fet3, inhibition of the recycling pathway results in targeting to the vacuole (Sato *et al.*, 2004), suggesting an additional layer of quality control operating in the late secretory pathway.

We have found that under normal conditions, a maturation-defective mutant of TREM2 is resistant to cleavage by endo D. Expression of nucleotide-locked forms of Arf1 to inhibit COPI-mediated recycling to the ER reveals endo D sensitivity for these mutants, suggesting that they normally reach a post-ER compartment *in vivo*. Although endo D sensitivity is classically used as a test for delivery of glycoproteins to the *cis*-Golgi compartment, endo D cleavage of VSV-G protein has been observed under conditions where it is trapped in the ERGIC (Alvarez *et al.*, 1999). In addition, α-mannosidase I, the key enzyme mediating sensitivity to endo D, can be detected in purified ERGIC membranes (Breuza *et al.*, 2004) and has been shown to cycle between the ERGIC and *cis*-Golgi (Alvarez *et al.*, 1999). In light of these findings and the data presented in this paper, it seems likely that the endo D sensitivity we observe for mutant TREM2 upon inhibition of the COPI pathway is a result of TREM2 becoming trapped in the ERGIC. One question that arises from our data is why mutant TREM2 does not accumulate in an endo D-sensitive form. One possibility is that mutant TREM2 is normally recycled from the ERGIC very efficiently, such that it cannot be fully processed by α-mannosidase I. That it is recycled efficiently is suggested by the observation of *O*-glycosylation as detected by interaction of a small percentage of the TREM2 molecules with the lectin jacalin, but not so efficiently glycosylated so as to be detected by a majority of molecules becoming sensitive to endo D. Consistent with this possibility, unassembled MHC molecules capable of reaching the Golgi remain insensitive to cleavage by endo D (Hsu *et al.*, 1991). This finding led to speculation that such MHC molecules may recycle to the ER from a very early Golgi compartment referred to as the *cis*-Golgi network (Hsu *et al.*, 1991), which is closely related or identical to the ERGIC (Glick, 2000; Klumperman, 2000). Alternatively, α-mannosidase I may become enriched in the ERGIC upon expression of nucleotide-locked Arf1 mutants, and this enrichment could account for the observed endo D sensitivity.

An additional question that arises from our work is why mutant TREM2 appears to become trapped in the ERGIC upon COPI knockdown, as opposed to moving further along the secretory pathway. Although COPI appears to be required for efficient retrieval of mutant TREM2 to the ER, COPI depletion apparently is not sufficient to enable mutant TREM2 to reach the later parts of the secretory pathway. This observation suggests an additional layer of quality control within the ERGIC, which could be mediated by an interaction between mutant TREM2 and a protein such as the KDEL receptor or Rer1. Such an interaction might be sufficient to prevent further anterograde transport of mutant TREM2 even when the retrograde pathway is blocked. An alternative possibility is that anterograde trafficking of mutant TREM2 further into the Golgi is inhibited as an indirect result of COPI depletion (Papanikou *et al.*, 2015).

Our findings raise the possibility that additional mutated and/or malformed membrane proteins that are prevented from reaching the cell surface and appear to be retained in the ER, are in fact transported to the ERGIC and subjected to ER retrieval. Although a role for an early Golgi cisterna in returning unassembled subunits of heteromeric membrane protein complexes has been established, our finding that disease-causing point mutations in a membrane protein induce cycling between the ER and ERGIC may represent a novel quality-control paradigm. In this context, it should be noted that under physiological conditions, TREM2 is normally found in complex with the signaling adaptor protein DAP12 at the cell surface (Colonna, 2003), and DAP12 is not expressed in the cell lines we have used for most experiments in this study. However, the lack of DAP12 is unlikely for several reasons to account for the recycling behavior we observe here. First, the recycling we observe is specific to mutant TREM2, and WT TREM2 reaches the cell surface efficiently in a variety of heterologous expression systems (shown here and in (Kleinberger *et al.*, 2014; Park *et al.*, 2015)). Second, co-expression of DAP12 with TREM2 (T66M) has been shown to be insufficient to alter this mutant’s trafficking phenotype (Kleinberger *et al.*, 2014). Third, we observe a striking correlation between defective maturation and *O*-glycan addition, underscoring the point that the cycling pathway we observe here is specific to particular TREM2 point mutants.

A better understanding of the ERGIC-to-ER quality-control pathway identified here will be of interest not only from a cell biological perspective, but also from a clinical perspective. Although homozygous inheritance of the mutations studied here is rare, heterozygous carriers may also be at increased risk for neurodegeneration (Guerreiro *et al.*, 2013a). For mutants that retain the ability to bind ligand, re-routing of the mutant protein to the cell surface by inhibiting recycling from and retention within the ERGIC could be a novel means of restoring TREM2-mediated signaling in cells that would otherwise be deficient. Finally, it will be interesting to determine whether recycling between the ER and ERGIC is a more widespread phenomenon than currently appreciated for pathogenic forms of membrane proteins, and if so, what signals and receptors, e.g. Rer1, are involved in recognizing mutant cargo for recycling.

## MATERIALS AND METHODS

### Antibodies

The HA.11 mAb used to detect HA-TREM2 by immunoblotting and cell-surface immunofluorescence was from Covance (Dedham, MA, USA); the CHC, EEA1 and GM130 mAbs were from BD Transduction Laboratories (San Jose, CA, USA); the ERGIC-53 mAb was from Enzo Life Sciences (Farmingdale, NY, USA); and the TfR mAb was from Invitrogen (Carlsbad, CA, USA). The TREM2 polyclonal Ab (pAb) used for three-color immunofluorescence and selected immunoblots was from R&D Systems (Minneapolis, MN, USA), the GFP pAb used to detect TREM2-GFP by immunoblot was from Torrey Pines Biolabs (Secaucus, NJ, USA), the calnexin pAb was from Abcam (Cambridge, MA, USA), the Sec31A pAb was from Bethyl Laboratories (Montgomery, TX, USA), and the LAMP2 pAb was from Sigma-Aldrich (St. Louis, MO, USA). The Sec22B and RPN I antisera were previously generated in our laboratory and described in (Schindler and Schekman, 2009). All Alexa Fluor-conjugated secondary antibodies used for immunofluorescence were from Life Technologies (Carlsbad, CA, USA).

### Molecular Biology

The human TREM2 cDNA was obtained from R&D Systems, amplified by PCR and inserted into the pEGFP-N1 vector after first removing the EGFP coding sequence. To facilitate detection of TREM2, we inserted an HA epitope tag and linker sequence (Kleinberger *et al.*, 2014) after the TREM2 signal sequence using the Phusion high-fidelity DNA polymerase system (NEB, Ipswich, MA, USA) for site-directed mutagenesis. To generate the HA-TREM2-GFP construct, we PCR amplified HATREM2 for insertion upstream of the EGFP coding sequence in pEGFP-N1. All TREM2 disease-associated missense mutations were also generated using Phusion-based mutagenesis, with the HA-TREM2 parent construct serving as template DNA. The *N-*glycosylation-defective version of HA-TREM2 (Y38C) had the following motifs in the TREM2 sequence mutated to alanine: ^20^NTT^22^ and ^79^NGS^81^. The Arf1 (T31N)-GFP construct was generated previously in our laboratory by Y. Guo. Arf1 (Q71L)-GFP was generated by site-directed mutagenesis of a WT Arf1-GFP construct. Constructs containing the synthetic mucin-type *O*-glycosylation motifs were generated based on a sequence (ATPAP) identified in (Yoneda *et al.*, 2001). In particular, an EcoRV site was first generated downstream of the N-terminal HA tag and glycine-rich linker sequence. Next, a DNA duplex encoding the (ATPAP)_8_ motif and additional C-terminal linker sequence was inserted into the EcoRV site. The resulting construct encoded HA-ATPAP)_8_-TREM2 (WT or Y38C). The version of TREM2 (Y38C) lacking its cytoplasmic tail was generated by PCR and encoded a stop codon after the glycine at position 200. All constructs were verified by sequencing at the UC Berkeley DNA Sequencing Facility.

### Cell Culture

Cell lines were maintained at the UC Berkeley Cell Culture Facility in a humidified chamber at 37°C and 5% CO_2_. CHO-Lec1 cells were obtained from ATCC (Manassas, VA, USA). All cell lines used in this study were grown in high-glucose DMEM supplemented with GlutaMAX (ThermoFisher, Waltham, MA, USA), 10% fetal bovine serum and 1% non-essential amino acids (NEAA), with the exception of U2OS cells, for which NEAA were not added. N9 cells were additionally supplemented with 1% sodium pyruvate. Cells were transiently transfected using Lipofectamine 2000 (ThermoFisher) according to the manufacturer’s instructions. Culture medium was typically changed 4 hr after transfection, and experiments were carried out the following day. N9 cells stably expressing HA-TREM2 and HA-TREM2-GFP (WT and Y38C) were generated by nucleofection (Lonza, Walkersville, MD, USA) using program A-30 and nucleofection kit V. To generate the stable transfectants, we nucleofected N9 cells with ApaLI-linearized plasmids and began selection with G418 (Life Technologies) at a concentration of 500 μg/ml two days post-nucleofection.

### Immunoblotting

Cells cultured in 6-well plates were harvested on ice by washing with 2 ml/well cold PBS followed by lysis in 150 μl of a buffer containing 100 mM NaCl, 10 mM Tris-Cl, pH 7.6, 1% (v/v) Triton X-100 and *Complete* protease inhibitor cocktail (Roche, Indianapolis, IN, USA). Triton-insoluble material was sedimented by centrifugation at 20,000*g* for 10 min at 4°C. Supernatants were mixed with 5X SDSPAGE sample buffer supplemented with DTT, then heated at 55°C for 10 min prior to running in 4-20% acrylamide gradient gels (Life Technologies and Bio-Rad, Hercules, CA, USA). After SDS-PAGE, proteins were transferred onto PVDF membranes (EMD Millipore, Billerica, MA, USA), blocked in 5% non-fat milk (dissolved in PBS containing 0.1% Tween-20), and probed with the following antibodies: HA.11 (1:2,500), CHC (1:10,000), TfR (1:10,000), TREM2 (1:1,000), Sec22B (1:10,000), RPN I (1:10,000), and GFP (1:5,000). Blots were developed using enhanced chemiluminescence and imaged on a ChemiDoc digital imager (Bio-Rad). Protein signals were quantified by densitometry using ImageJ (NIH, Bethesda, MD, USA).

### Cell Surface Biotinylation

Cell surface biotinylation was carried out as described in (Sirkis *et al.*, 2016). Cells cultured in 6-well plates were washed twice with 2 ml/well room temperature (RT) PBS and labeled with 1 ml/well EZ-Link Sulfo-NHS-SS-Biotin reagent (ThermoFisher) at 1 mg/ml in PBS for 15 min. Cells were then placed on ice, washed with 2 ml/well cold Tris-buffered saline to quench the biotin reagent, then washed with 2 ml/well cold PBS and finally lysed and clarified as described above. To capture biotinylated proteins, we added 7.5 μl *Strep*-Tactin resin (IBA Lifesciences, Göttingen, Germany) to 100 μl clarified lysates and the mixtures were rotated at 4°C for 1 hr. The resin was then sedimented and washed multiple times with lysis buffer. Finally, 2X SDS-PAGE sample buffer containing DTT was added to the washed resin, the samples were mixed with the use of a vortex, then heated and processed for immunoblotting as described above.

### *In vitro* Budding Assays

The budding assay was performed essentially as described previously (Kim *et al.*, 2007; Merte *et al.*, 2010), with modifications described below. Briefly, one 10-cm plate of HEK-293T cells was used per transfection condition. To harvest cells for the budding assay, we first washed cells with 10 ml cold PBS, then lifted them from the plate by gentle pipetting with 10 ml cold KHM buffer (110 mM KOAc, 2 mM MgOAc, 20 mM HEPES, pH 7.2). Cells were sedimented at 300*g* for 5 min, resuspended in 6 ml cold KHM and gently permeabilized during a 5 min incubation on ice with 20 μg/ml digitonin, then washed during a 5 min incubation on ice with high-salt KHM (containing 300 mM KOAc). The reaction (100 μl) was started by incubating semi-intact cells (at an OD_600_ of 1.0) at 30°C in KHM supplemented with 3 mg/ml rat liver cytosol, an ATP regeneration system, and 200 μM GTP or GTPγS. Rat liver Cytosol was prepared as described in (Kim *et al.*, 2005). Where indicated, Sar1A (H79G) (purified as in (Kim *et al.*, 2005)) was added at a concentration of 10 μg/ml. For budding assays with N9 cell membranes, digitonin was used at a concentration of 40 μg/ml to achieve optimal permeabilization. Budding efficiency was determined by comparing the immunoblot signal intensity (measured as described above) of budded lanes to that of the appropriate 5% membrane input lane.

### Endoglycosidase Digests

Endo D, endo H and PNGase F were obtained from NEB. Endo F1, F2 and F3 were obtained from Sigma-Aldrich. Digests were performed on cell lysates (prepared as described above) according to the manufacturers’ recommendations. Digests were carried out at 37°C for 1 hr and included 2 μl of the appropriate enzyme in 20-μl reactions, with the following exceptions: for PNGase F, 1 μl of enzyme was used, and for the endo F enzymes, 50-μl reactions were used. For digests with endo D, lysates were supplemented with DTT and heated at 95°C for 3 min prior to starting the digest. For digests with endo H and PNGase F, lysates were supplemented with glycoprotein denaturing buffer and heated at 95°C for 5 min prior to starting the digest. For digests with endo F enzymes, lysates were not denatured prior to beginning the reaction. Reactions were stopped by the addition of 5X SDS-PAGE sample buffer containing DTT, followed by heating at 65°C for 10 min.

### Jacalin Affinity Chromatography

Cell lysates were prepared as described above for immunoblotting. Agarose-bound jacalin (Vector Labs, Burlingame, CA, USA) was pre-equilibrated in cell lysis buffer and 20 μl of resin was added to 900 μl of clarified cell lysates. Mixtures of lysate and resin were rotated at 4°C for 1 hr. The resin was then centrifuged and washed multiple times with lysis buffer. Finally, 2X SDS-PAGE sample buffer supplemented with DTT was added to the washed resin, the samples were dispersed with the use of a vortex mixer, then heated and processed for SDS-PAGE and immunoblotting as described above.

### siRNA Transfection

We obtained four distinct siRNAs targeting δ COP (*ARCN1* gene) (FlexiTube GeneSolution; Qiagen, Germantown, MD, USA). siRNAs #4 (target sequence: AAGGCTGAGATGCGTCGTAAA) and #7 (target sequence: AGGCAACTGATTGTTTCGATA) were selected based on their knockdown efficiency and lack of toxicity. U2OS cells were transfected with these siRNAs or a non-targeting control siRNA at a final concentration of 20 nM using the Lipofectamine RNAiMAX reagent (Life Technologies) according to the manufacturer’s instructions. Cells were typically transfected with the appropriate plasmid for immunofluorescence imaging on the following day, and cells were processed for microscopy (see below) approximately 48 h after the siRNA transfection.

### Immunofluorescence Microscopy

For cell surface labeling of HA-TREM2, we fixed U2OS cells by adding an equal volume of 4% EM-grade formaldehyde (Electron Microscopy Sciences, Hatfield, PA, USA) diluted in PBS to the cell culture medium and incubating for 20 min at RT. Cells were then washed 3X with PBS and blocked without permeabilization using a buffer containing 2% bovine serum albumin and 1% fish skin gelatin in PBS. Cell surface HA was detected using the HA.11 mAb diluted 1:500 in blocking buffer and incubating for 1 h at RT. Cells were washed as above and incubated for 45 min at RT with an anti-mouse IgG secondary Ab conjugated to Alexa Fluor 568 diluted 1:500 in blocking buffer. Cells were again washed as above, rinsed briefly in dH2O, then mounted on slides using ProLong Gold antifade reagent containing DAPI (ThermoFisher). Fluorescence from the cytoplasmic GFP tag on TREM2 was detected directly.

For the other immunofluorescence experiments a similar procedure was followed, except that the blocking buffer also contained either 0.1% Triton X-100 (for ERGIC-53 labeling) or 0.02% saponin (for all other antibodies) to permeabilize the cells after fixation. In these experiments, the calnexin pAb was used at 1:200, the ERGIC-53 mAb at 1:400, the Sec31A pAb at 1:200, the GM130 mAb at 1:200, the EEA1 mAb at 1:200, and the LAMP2 pAb at 1:500. For three-color immunofluorescence imaging involving Arf1 (Q71L)-GFP, untagged TREM2 was detected using the TREM2 pAb at 1:200. GFP fluorescence was detected directly and marker proteins were detected using the appropriate secondary Ab conjugated to Alexa Fluor 568 (1:1,000), with the exception of the TREM2 pAb, which was detected using an anti-goat IgG secondary Ab conjugated to Alexa Fluor 647 (1:1,000). Cells were imaged using an inverted epifluorescence microscope (Axio Observer.Z1, Zeiss, Thornwood, NY, USA) and a Plan-Apochromat 63x, 1.4 NA objective lens. Tiff files were analyzed, cropped and pseudo-colored in ImageJ (NIH).

## ACKNOWLEDGMENTS

The authors thank J. Brodsky, A. Helenius, J. Olzmann, V. Malhotra, C. Asensio and members of the Schekman lab for helpful comments and questions. We thank the UC Berkeley Cell Culture Facility for maintenance of cell lines and the UC Berkeley DNA Sequencing Facility. D.S. is funded by NIH fellowship F32 AG050404 and was previously funded by HHMI. R.A. is funded by HHMI and was previously funded by a SURF/Rose Hills summer fellowship. R.S. is funded by HHMI.

## FIGURES AND FIGURE LEGENDS

**Supplemental Figure S1.**
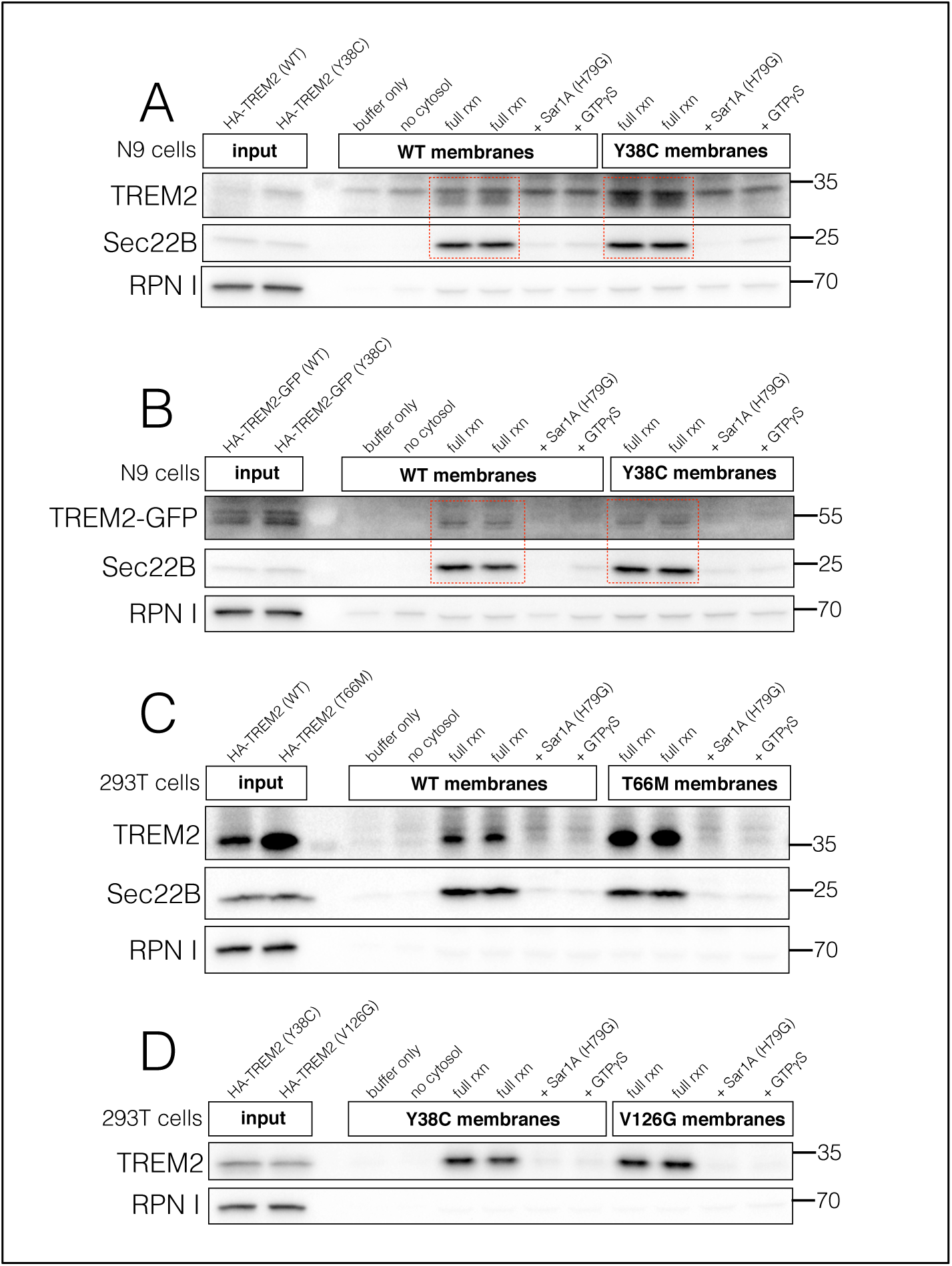
*In vitro* budding of TREM2 (Y38C) from N9 cell membranes and budding behavior of additional TREM2 mutants. Membranes from N9 cells stably expressing WT or Y38C HA-TREM2 (A) or HA-TREM2-GFP (B) were used in *in vitro* budding assays. HA-TREM2 (Y38C) and HA-TREM2-GFP (Y38C) show budding behavior similar to the WT protein. A non-specific band running at ~35 kD (A) migrates just above the specific HA-TREM2 band. Budding assays performed using membranes from HEK-293T cells transiently expressing HA-TREM2 (T66M) (C) and (V126G) (D) demonstrate that additional TREM2 point mutants bud efficiently from the ER.

**Supplemental Figure S2.**
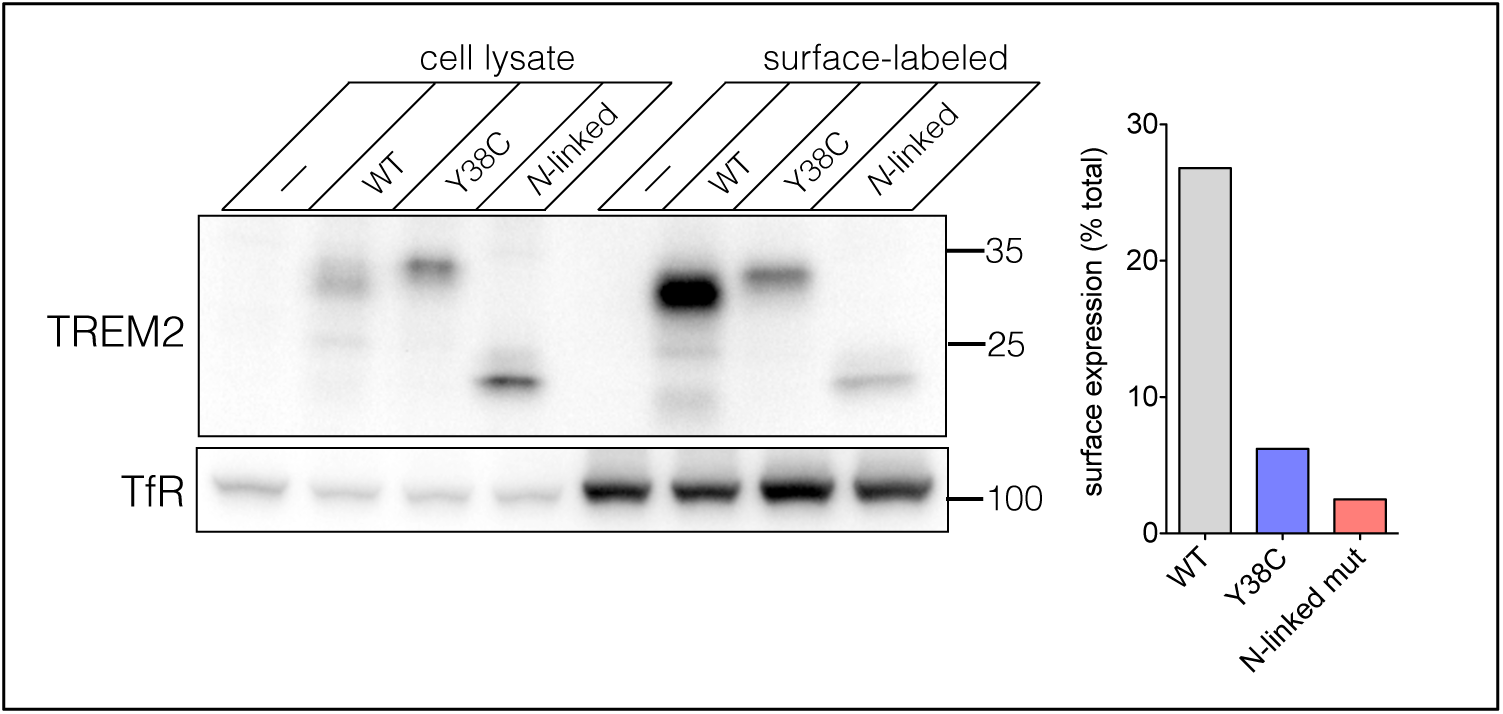
Cell surface biotinylation of WT and mutant TREM2 in CHO-Lec1 cells. Surface biotinylation was performed as in Figure 1. HA-TREM2 (Y38C) surface expression is impaired relative to WT in CHO-Lec1 cells. A version of the Y38C mutant rendered defective for *N*-linked glycosylation was used as a control protein for lack of surface expression. TfR was used as a positive control for surface expression and as a loading control.

**Supplemental Figure S3.**
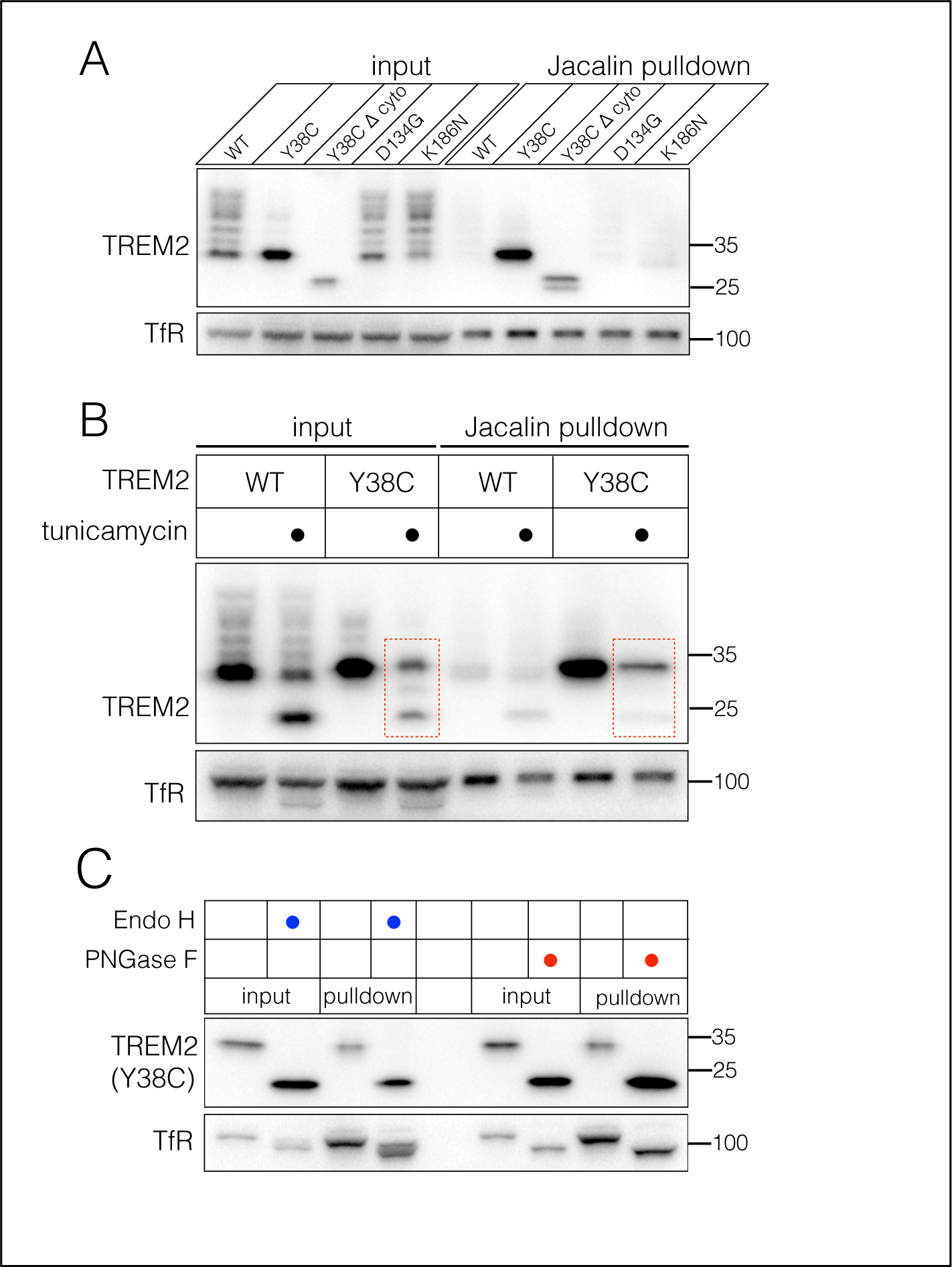
Control experiments for jacalin affinity chromatography. (A) The cytoplasmic tail of TREM2 is not required for pulldown of the Y38C mutant by jacalin. Mutants D134G and K186N, which do not display maturation defects, are not bound by jacalin. (B) A non-glycosylated form of TREM2 (Y38C) induced by tunicamycin treatment is not efficiently pulled down by jacalin, while a glycosylated form derived from the same lysates is pulled down (highlighted by dashed red rectangles). (C) Enzymatic removal of *N*-linked glycans from TREM2 (Y38C) by endo H or PNGase F does not affect pulldown by jacalin.

**Supplemental Figure S4.**
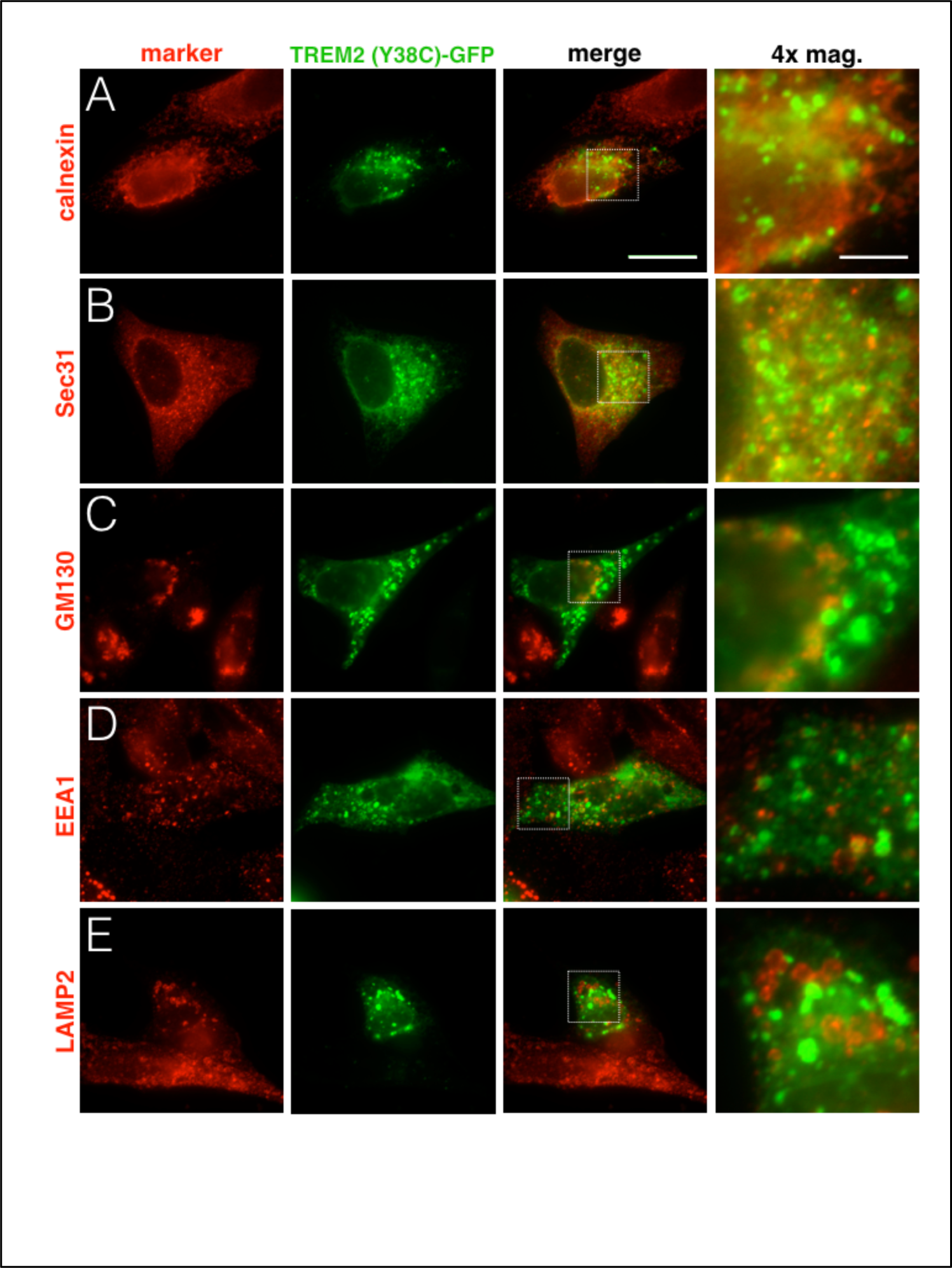
Additional organelle markers that do not colocalize with TREM2 (Y38C) after COPI knockdown. U2OS cells were transfected with siRNA against δ COP and processed for immunofluorescence as in Figure 5. TREM2 (Y38C)- GFP puncta do not show appreciable colocalization with calnexin, Sec31, GM130, EEA1 or LAMP2. Scale bars indicate 20 μm for standard magnification and 5 μm for the 4x-magnified images.

